# Spectroscopy-Based Machine Learning to Predict Stem-End Rot in Avocados with the Temporal Effects

**DOI:** 10.64898/2026.10.29.762723

**Authors:** Xuezhen Guo, Aneesh Chauhan

**Affiliations:** Wageningen Food & Biobased Research, the Netherlands

**Keywords:** non-destructive, Fruit quality, Artificial intelligence, Neural Networks, Vis-NIR, Postharvest, Temporal Effects

## Abstract

Accurate prediction of postharvest fruit quality is essential for reducing food waste and improving supply chain efficiency. This study investigates the integration of spectral information and temporal dynamics for predicting stem-end rot in avocados using hyperspectral Vis-NIR data. Three modelling approaches were compared: a feedforward neural network using only spectral information, an extended model incorporating storage duration, and a physics-informed neural network (PINN) encoding a first-order kinetic constraint. To evaluate temporal extrapolation, models were trained on samples from early storage stages and tested on day-12 data. Results show that while spectral data provide strong predictive capability, incorporating time as a simple input can lead to overfitting. In contrast, the PINN improves generalization by guiding learning with mechanistic knowledge, achieving the best performance on unseen temporal conditions. These findings highlight the value of combining data-driven and physics-based approaches, offering a promising pathway toward more robust and interpretable models for time-dependent food quality assessment.

## 1. Introduction

Fresh produce, such as fruits and vegetables, undergoes a natural quality decay process, leading to substantial food waste in postharvest supply chains. Accurately predicting the quality of fresh produce is crucial for businesses aiming to keep competitive in this sector. As a result, great efforts have been dedicated to developing accurate models for predicting the quality of fresh produce (Chauhan et al., 2023; Wang HaiLong et al., 2015).

The degradation in quality of fresh produce is largely driven by a series of biochemical reactions occurring within the products, which are regulated by kinetic processes. These reactions follow the Arrhenius law, a principle that explains how reaction rates depend on temperature. Therefore, temperature and the duration of exposure to these conditions are the primary factors that influence the rate of quality decay in fresh produce. Due to this relationship, the most widely used models for predicting quality decay are mechanistic (kinetic) models. These models are based on the relationship between the quality indicator and time-temperature dynamics and are often represented by a logistic curve (Tijskens & Polderdijk, 1996; Tromp et al., 2016; Tromp et al., 2017; Tromp et al., 2012). A key strength of classical kinetic models lies in their biological foundation, as demonstrated by Tijskens and Polderdijk (1996). These models are grounded in product physiological knowledge, which makes them both powerful and highly explainable. As such, they are widely regarded as effective knowledge-based modelling approaches. Parallel to knowledge-based approaches, data-driven approaches have also been developed. For example, Lammertyn et al. (2000) employed multinomial logistic regression to predict the quality of conference pears, allowing for the inclusion of additional variables to capture the variability within and between different product batches. Zhu et al. (2017) conducted a study in which they developed a fuzzy Bayesian network model to evaluate and control the quality of fresh food in O2O e-commerce, identifying critical factors along the supply chain and determining key intervention points such as unloading, warehouse inspection, and monitoring. Singh et al. (2009) used artificial neural networks to predict the sensory quality of UHT-treated milk during storage based on physicochemical indicators of deteriorative reactions, and compared the prediction performance with traditional kinetic (mechanistic) models, showing that the ANN approach achieved higher accuracy. Within the data-driven modelling stream of literature, a prominent body of work focuses on models that utilize near-infrared spectroscopy (NIRS) to measure product quality. NIRS is widely applied because it enables rapid and non-destructive assessment of internal quality attributes of fresh produce. For example, Lamptey et al. (2023) investigated the use of a handheld NIR spectrometer combined with multivariate algorithms to identify mango varieties and predict key quality indicators such as total soluble solids (TSS) and pH. Their results demonstrated that NIR spectroscopy coupled with appropriate chemometric models can provide rapid and non-destructive assessment of fruit quality. Similarly, Téllez-García et al. (2025) evaluated portable NIRS sensors for the in situ assessment of physicochemical quality and maturity of intact mangoes. Their results showed that NIRS can accurately predict quality attributes such as soluble solids content and dry matter to support optimal harvest timing, although pre-calibrated commercial models may not always generalize well across different cultivars and growing conditions. In another study, Mancini et al. (2020) investigated the application of near-infrared spectroscopy combined with chemometric techniques to evaluate quality parameters of strawberry fruits. Their results showed that models based on principal component analysis and partial least squares regression could successfully predict key attributes such as soluble solids content and firmness, demonstrating the potential of NIR as a non-destructive alternative to conventional laboratory measurements. More recently, Amoriello et al. (2025) explored the use of portable Vis-NIR spectroscopy and hyperspectral imaging combined with machine learning techniques to monitor internal quality attributes of apricot fruits. Their findings indicated that artificial neural network models could accurately predict important quality parameters such as total soluble solids and dry matter, highlighting the capability of spectral sensing technologies for efficient quality monitoring. Furthermore, Elsayed et al. (2025) investigated the use of hyperspectral reflectance indices combined with machine learning models to estimate biochemical quality parameters of mango and strawberry fruits at different ripening stages. Their results demonstrated that machine learning models, including artificial neural networks, random forests, and decision trees, can accurately predict attributes such as chlorophyll content, total soluble solids, and firmness, illustrating the effectiveness of hyperspectral sensing for rapid fruit quality evaluation. Collectively, these studies demonstrate that NIR and hyperspectral sensing technologies, particularly when combined with chemometric and machine learning approaches, provide powerful tools for rapid and non-destructive assessment of fruit quality across different crops and supply chain stages. Chauhan et al. (2024) conducted a study using hyperspectral NIR data to predict avocado stem-end rot after storage, applying various machine learning approaches including Random Forest, XGBoost, and semi-supervised deep autoencoder classifiers. While promising, their study did not consider fruit maturity, age, or other time-dependent physiological changes, even though disease progression is likely influenced by temporal processes during ripening and storage. Building on this research, the present study investigates the incorporation of temporal effects into NIR-based models to assess whether this integration can improve predictive accuracy, with a particular focus on enhancing the model’s extrapolation capability over time during storage.

For quality monitoring, an emerging direction is the use of hybrid modelling frameworks that combine kinetic or mechanistic models with data-driven approaches to leverage both domain knowledge and empirical data. For example, Xiao and Li (2022) integrated the closed-form solution of a kinetic model with smoothed near-infrared spectral data using functional regression to predict the firmness of fresh produce. More recently, van der Sman et al. (2026) and Guo et al. (2026) employed scientific machine learning approaches that combine neural networks with mechanistic equations to predict the quality evolution of fruit and vegetable products during storage. Compared with purely data-driven approaches, these hybrid models provide the additional advantage of embedding physical interpretability and biological realism into the learning process. By constraining machine learning models with known physical laws or kinetic relationships, scientific machine learning can improve model robustness, reduce overfitting, and enhance extrapolation capability under unseen conditions. This is particularly important for postharvest quality prediction, where product deterioration is inherently a dynamic and time-dependent process. By combining the biological foundation of mechanistic models with the flexibility and representation power of machine learning, hybrid modelling approaches have shown strong potential for improving predictive accuracy while also providing more interpretable and scientifically meaningful predictions. Inspired by these developments, the present study explores whether incorporating temporal kinetic knowledge into hyperspectral learning models can improve the prediction and temporal extrapolation of avocado stem-end rot during storage.

## 2. Materials and Methods

### 2.1. Data collection

The avocados used in this study were imported from the Huantar region of Peru during May and June 2018 and subjected to a natural infection process. The experimental assessment was conducted by researchers at Wageningen Food & Biobased Research. The experimental design is illustrated in Figure 1. A total of 236 avocados with similar initial firmness levels were selected to ensure a relatively homogeneous starting condition, thereby minimizing variability unrelated to the infection process. The fruits were stored under controlled conditions that allowed the natural development of stem-end rot without artificial inoculation, ensuring that the observed infection patterns reflect realistic postharvest scenarios.

**Figure 1.**
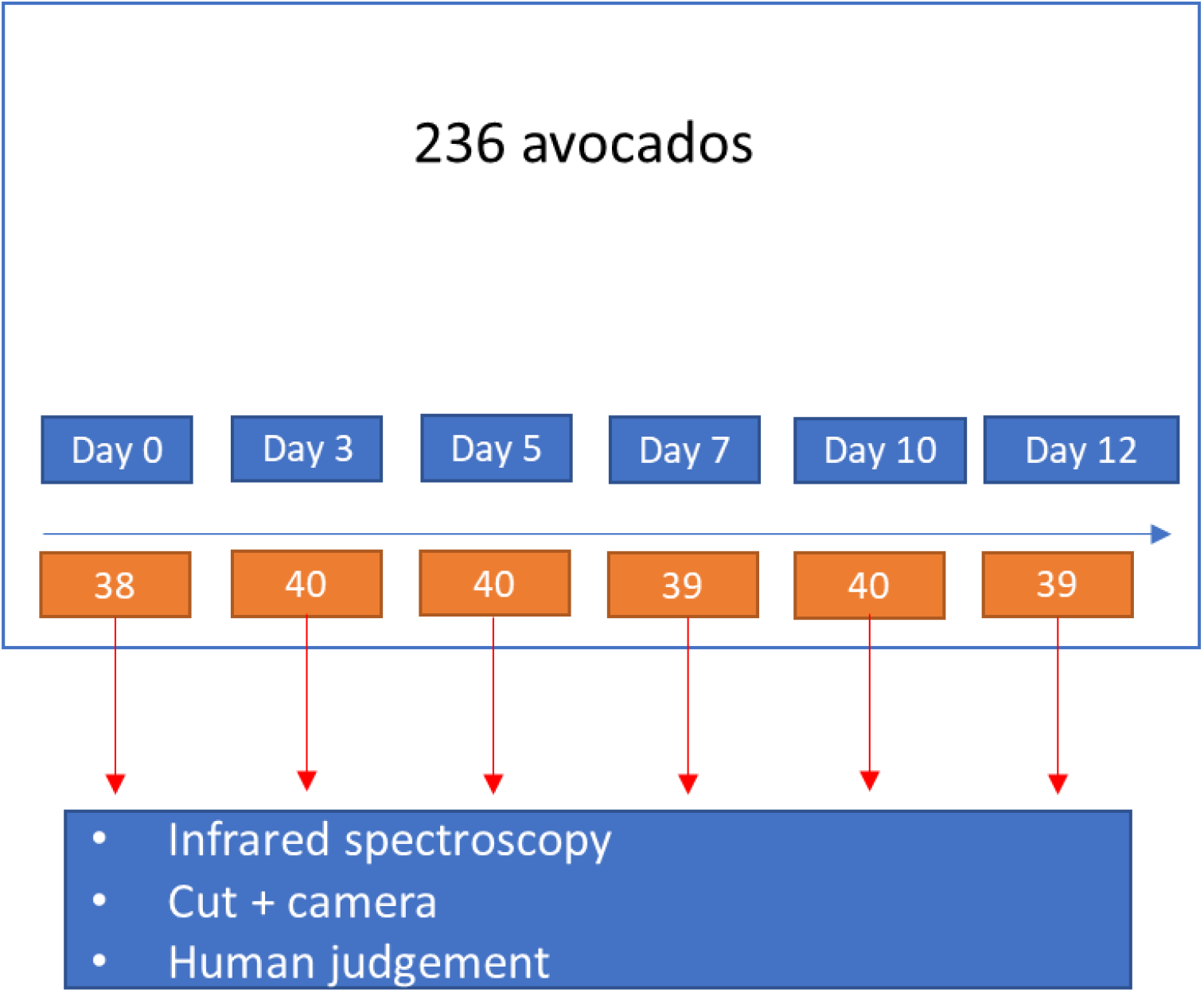
The experimental design

On each specified measurement day (i.e., after a defined storage duration)—day 0, 3, 5, 7, 10, and 12—approximately 40 avocados were randomly selected from the batch. These samples were destructively evaluated by cutting them open and assessing the presence and severity of stem-end rot. The assessment was conducted by trained experts using both direct visual inspection and camera imaging, providing a reliable and consistent labelling of the infection status. This procedure ensured that the ground-truth labels used for model development accurately reflect the internal quality condition of each fruit. By distributing the measurements across multiple time points, the dataset captures the progressive nature of the infection process and provides a temporal dimension that is critical for subsequent modelling.

Simultaneously with the destructive assessment, hyperspectral images were acquired for each individual avocado using a portable visible–near-infrared (Vis–NIR) spectral camera (SpecimIQ, Specim, Finland). This device enables non-destructive measurement of spectral reflectance, capturing both spatial and spectral information from the fruit surface. The camera recorded reflectance values across the wavelength range of 400 to 1000 nm, covering both the visible region associated with pigment-related characteristics and the near-infrared region linked to internal chemical composition. The spectral sampling interval of 7 nm resulted in a high-dimensional representation of each sample, allowing subtle variations in spectral signatures to be captured.

The spectra of all 236 avocados are presented in Figure 2. These spectra provide an overview of the variability present in the dataset, reflecting differences in fruit composition, maturity, and infection status. While individual spectra exhibit similar overall patterns, noticeable variations can be observed across samples, particularly in specific wavelength regions associated with water content, sugars, and structural components. Such variability forms the basis for developing predictive models that link spectral features to internal quality attributes.

**Figure 2.**
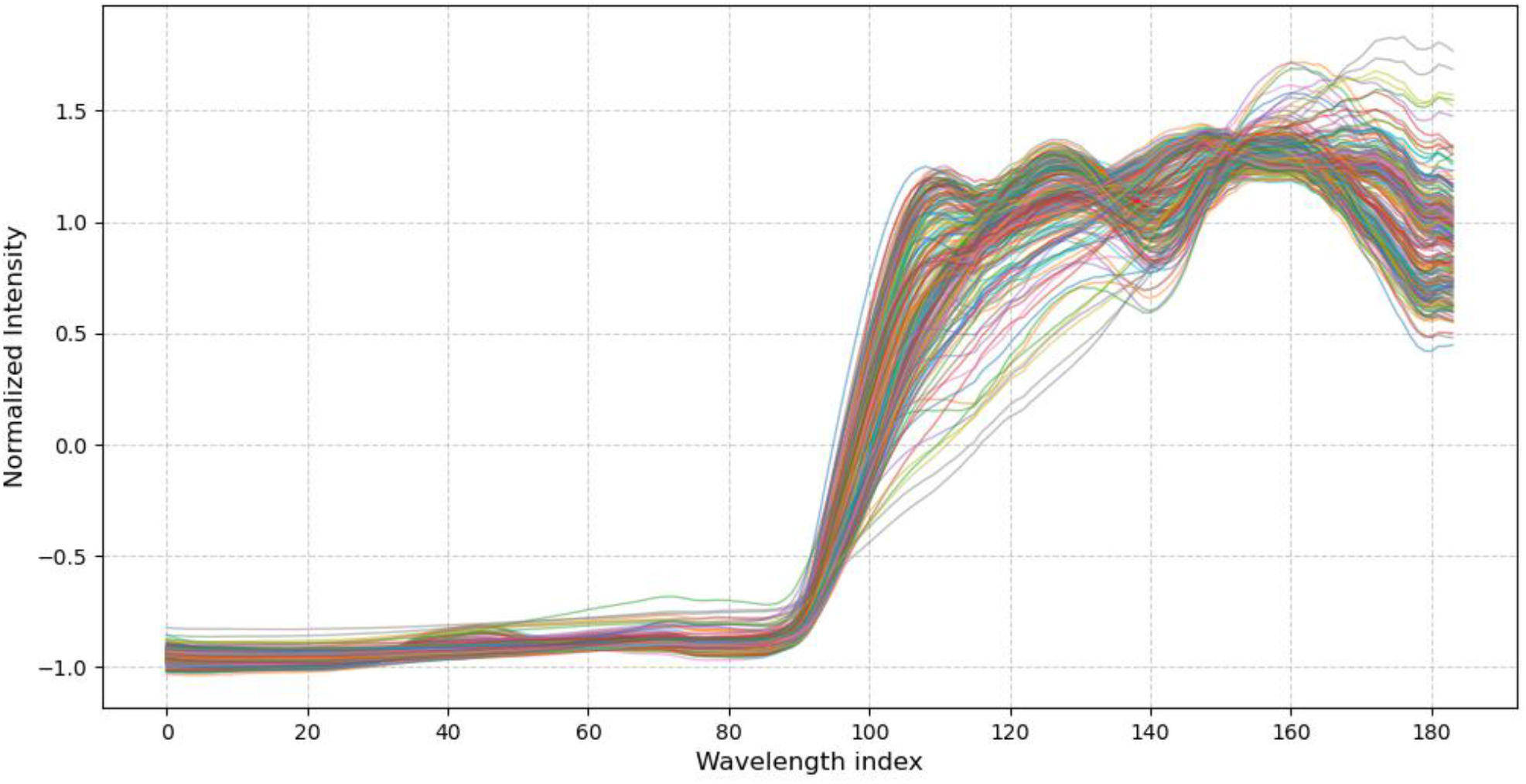
The spectra of all 236 avocadoes

Figure 3 further illustrates the spectral profiles of avocados grouped by storage duration. Compared to the aggregated spectra shown in Figure 2, this representation highlights how spectral characteristics evolve over time. It can be observed that, as storage time increases, the shape and intensity of the spectra exhibit a systematic drift. In particular, shifts in reflectance levels and subtle changes in spectral curvature suggest underlying biochemical and structural transformations occurring during storage and infection progression. These changes may be associated with moisture loss, tissue degradation, and metabolic activity linked to fungal growth.

**Figure 3.**
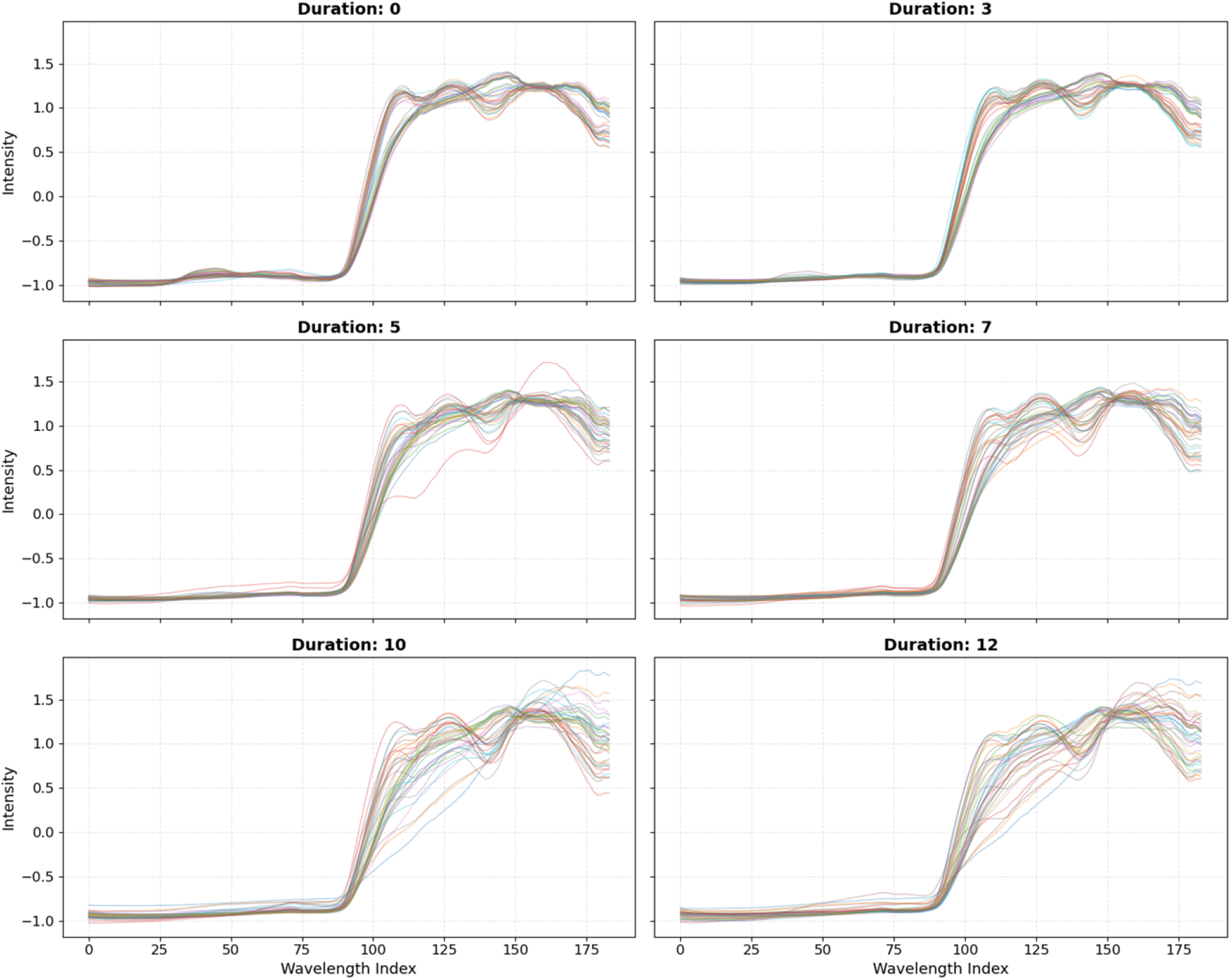
The spectra of all 236 avocadoes by storage duration

Importantly, the observed temporal evolution of the spectra provides initial evidence that time-dependent information is embedded within the spectral data. This suggests that incorporating temporal features, or explicitly modelling temporal dynamics, may enhance the ability to predict stem-end rot. The gradual and consistent nature of the spectral shifts indicates that the infection process is not abrupt but follows a continuous progression, which can potentially be captured using appropriate modelling techniques.

Table 1 presents the distribution of the binary target variable, with counts of 0 and 1 representing healthy and infected (stem-end rot) avocados, respectively. As expected, the distribution exhibits a clear temporal pattern, with the proportion of infected samples increasing over storage time.

**Table 1.** The distribution of the healthy and infected avocadoes after different storage durations.

| Duration (Days) | Healthy avocado (0-count) | Infected avocado (1-count) | Infected avocado (%) |
| --- | --- | --- | --- |
| 0 | 38 | 0 | 0.00% |
| 3 | 40 | 0 | 0.00% |
| 5 | 38 | 2 | 5.00% |
| 7 | 36 | 3 | 7.70% |
| 10 | 32 | 8 | 20.00% |
| 12 | 27 | 12 | 30.80% |
| **Total** | **211** | **25** | **10.60%** |

Table 1 presents the distribution of the binary target variable, where values of 0 and 1 represent healthy and infected (stem-end rot) avocados, respectively. The table summarizes the number of samples in each class across different storage durations, providing insight into the evolution of infection prevalence over time. As expected, the distribution exhibits a clear temporal pattern: at early time points (e.g., day 0 and day 3), the majority of samples are classified as healthy, while at later stages (e.g., day 10 and day 12), the proportion of infected samples increases substantially.

This temporal imbalance reflects the natural progression of stem-end rot and highlights the increasing difficulty of maintaining fruit quality during extended storage. From a modelling perspective, this evolving class distribution introduces additional challenges, such as potential class imbalance and shifting data distributions across time. At the same time, it may imply the importance of incorporating temporal information into predictive models, as the probability of infection is inherently linked to storage duration.

### 2.2. Modelling approach

To systematically investigate the temporal effects of spectral data on predicting stem-end rot in avocados, three modelling approaches with increasing levels of complexity and physical interpretability were implemented. These include: (1) a feedforward neural network (FNN) using only the mean spectral values as input features (FNN_mean_only), (2) an FNN incorporating both mean spectral values and storage duration as input features (FNN), and (3) a physics-informed neural network (PINN) that integrates a first-order kinetic constraint into the learning objective, thereby enabling a scientific machine learning (SciML) framework. The progression from the first to the third model reflects a transition from purely data-driven learning toward hybrid modelling that combines empirical data with domain-specific knowledge.

The primary motivation for this comparative framework is to explicitly evaluate the role of temporal information in predicting the progression of stem-end rot. While hyperspectral data provide rich biochemical and structural information, the infection process itself evolves dynamically over time. Therefore, incorporating temporal features—or even more fundamentally, embedding known temporal dynamics—has the potential to significantly enhance predictive performance, particularly when extrapolating beyond the observed time range.

In the baseline FNN_mean_only model, only the mean reflectance of the hyperspectral signal is used as input. This model effectively ignores temporal information and serves as a reference to assess how much predictive power is contained in spectral features alone. The second model (FNN) augments this representation by including storage duration as an additional input variable. This allows the model to learn temporal patterns directly from the data in a purely empirical manner, without imposing any structural assumptions about the underlying dynamics.

The third model (PINN) further extends this idea by incorporating prior knowledge about the infection process through a physics-informed constraint. Specifically, the development of stem-end rot is assumed to follow first-order kinetic behaviour, which is commonly used to describe biological degradation and growth processes. The governing equation for a first-order kinetic model can be written as:

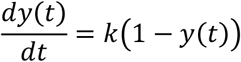

where

*y*(*t*) represents probability of infection at time *t*

*k* is a rate constant that determines how rapidly the infection progresses.

According to Boyce et al. (2009), the solution to this differential equation is given by:

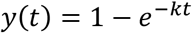

Within the PINN framework, this kinetic relationship is incorporated into the training process as a soft constraint rather than a hard requirement. This is achieved by augmenting the standard data-driven loss function with an additional physics-based loss term that penalizes deviations from the expected kinetic behaviour.

Specifically, the model is trained using a composite loss function consisting of a data-driven term and a physics-based term. The data-driven component is defined using the binary cross-entropy loss:

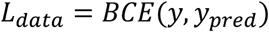

where *y* is the ground truth (0 or 1) and *y*_*pred*_ is the value (between 0 and 1) predicted by the PINN model.

To enforce the first-order kinetic behaviour, a residual term is constructed based on the governing differential equation:

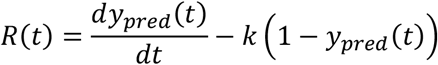

This physics-based loss is then defined as the mean squared error of this residual:

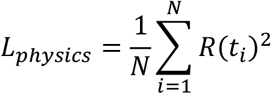

The total loss is therefore:

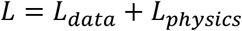

The trainable parameter *k* allows the PINN model to learn this parameter directly from the data. By doing so, the PINN not only predicts infection outcomes but also captures an interpretable parameter governing the infection dynamics.

All three models were trained using samples collected from earlier time points (days 0 to 11), while observations from day 12 were reserved as an independent test set. This experimental design is particularly important for evaluating temporal extrapolation, as it requires the models to generalize beyond the time range encountered during training. Such a setup closely mimics real-world scenarios in postharvest monitoring, where predictions are often needed for future storage conditions that have not yet been observed. Particular emphasis was therefore placed on evaluating model performance on the day-12 test set, as this provides a direct measure of each model’s ability to capture temporal dynamics and generalize to unseen future conditions.

To ensure a fair comparison, the neural network architectures were kept consistent across all models. Each model consists of two fully connected hidden layers with 32 neurons per layer and ReLU activation functions, followed by an output layer with a sigmoid activation function that produces a probability score between 0 and 1, corresponding to the likelihood of infection. Training was conducted using the Adam optimization algorithm with a learning rate of 0.001 over 2000 epochs. To monitor performance and prevent overfitting, the training data were further split into training and validation subsets using an 80:20 ratio, and the validation loss was tracked throughout the training process to ensure stable convergence.

To assess model performance, predicted probabilities were converted into binary class labels using the standard threshold of 0.5. Based on the comparison between predicted and true labels, each prediction was categorized as a true positive, true negative, false positive, or false negative. These outcomes were used to construct confusion matrices and derive classification metrics that quantify predictive accuracy. The comparison between the three models enables a clear assessment of (i) the contribution of spectral information alone, (ii) the added value of explicitly including time as an input feature, and (iii) the benefit of embedding mechanistic knowledge through physics-informed learning.

Finally, in addition to using the day-12 samples as an independent test set, we also evaluated the models using a traditional 80%–20% random train–test split for comparison, while keeping all other model settings unchanged.

## 3. Results and discussion

### 3.1. Results using day-12 samples as the testing Dataset

This section presents the results obtained using the day-12 samples as the testing dataset and the remaining samples as the training dataset. Table 2 reports the overall prediction accuracy of the three models on both the training and testing datasets. All models achieved strong performance on the training dataset, with the FNN_mean_only model yielding the highest accuracy (0.954), slightly outperforming the other two models. As expected, the prediction accuracy of all three models decreased on the testing dataset, since the day-12 samples should have patterns that differ from those observed during training. However, an interesting finding is that incorporating time as an additional feature led to a notable reduction in testing accuracy for the FNN_mean_time model, suggesting potential overfitting to the training data. In contrast, the inclusion of the physics-based loss in the PINN model resulted in a slight improvement in testing accuracy, although the gain is marginal (even neglectable). Notably, when considering both training and testing performance, the PINN model exhibited the lowest training accuracy but the highest testing accuracy among the three models. This pattern indicates that incorporating the physical constraint into the loss function may act as a form of regularization, which helps to mitigate overfitting and improve generalization to unseen temporal conditions.

**Table 2.** Overall model prediction accuracy on the training dataset and the day-12 testing dataset.

| Model | Training accuracy | Testing accuracy |
| --- | --- | --- |
| FNN_mean_only | 0.954 | 0.821 |
| FNN_mean_time | 0.944 | 0.667 |
| PINN | 0.939 | 0.846 |

A closer examination of the confusion matrices provides a more detailed view of classification performance. The FNN_mean_only model correctly identified 4 and 5 infected avocados in the training and testing datasets, respectively, without producing any false positives (Figure 4). This behaviour can be attributed to the fact that the training dataset is dominated by healthy avocados. As a result, the model is highly effective at detecting healthy samples but adopts a conservative approach when predicting infection, only classifying an avocado as infected when its spectral patterns are markedly different from those of healthy samples.

**Figure 4.**
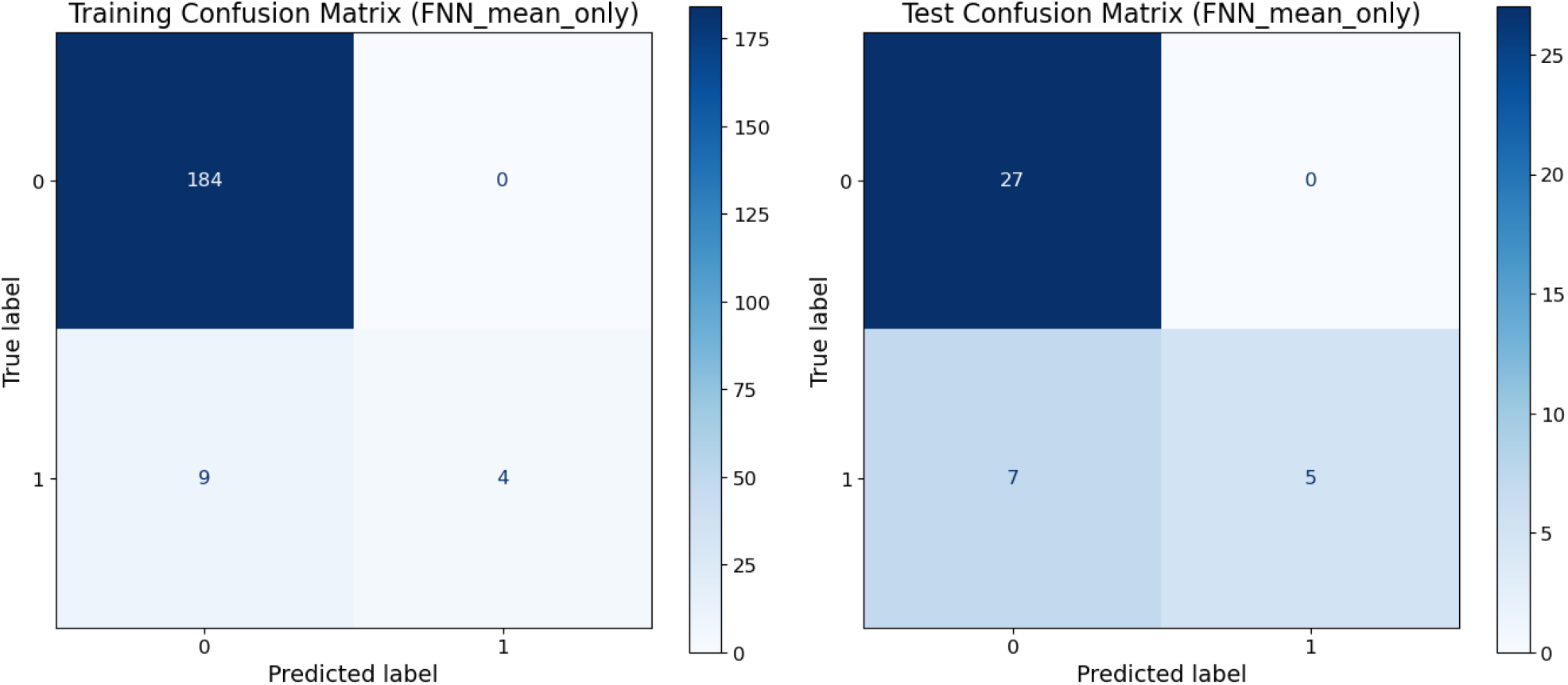
The confusion matrices for the FNN_mean_only model on the training dataset and the day-12 testing dataset

In contrast, the FNN_mean_time model correctly identifies a higher number of infected avocados—7 in the training set and 9 in the testing set—but this comes at the cost of substantially more false positives, with 5 and 10 misclassifications in the training and testing datasets, respectively (Figure 5). These results suggest that simply including time as an additional feature in a purely data-driven manner may disrupt the model’s ability to fully leverage the spectral information. The model appears to overemphasize temporal effects, effectively overfitting to the time dimension, leading it to predict infection more often than it should be.

**Figure 5.**
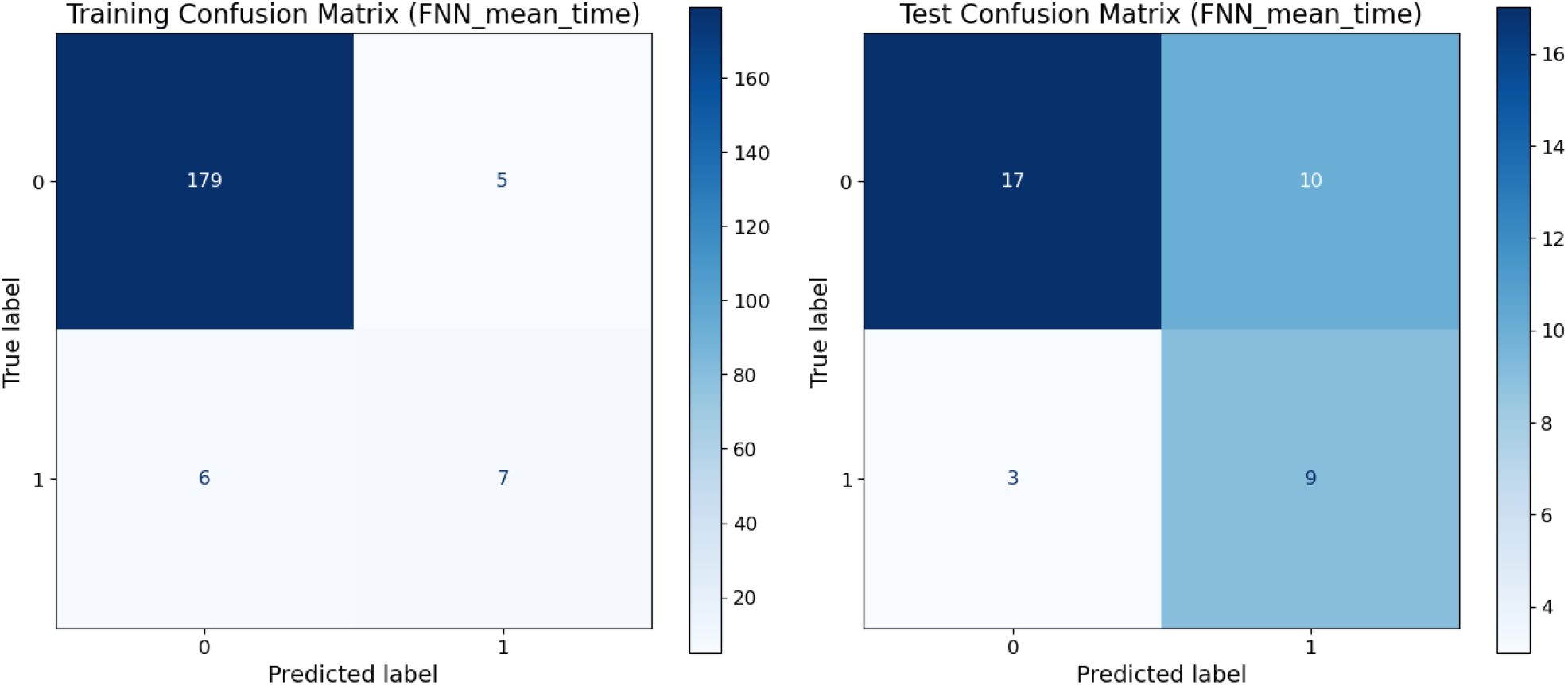
The confusion matrices for the FNN_mean_time model on the training dataset and the day-12 testing dataset

The overfitting to the time dimension appears to be substantially mitigated by the physics-informed loss component in the PINN. As shown in Figure 6, the correct classification of infected avocadoes is 8 on the testing dataset (compared to 9 for FNN_mean_time model), resulting in only 2 false positives (versus 10 for FNN_mean_time model). This suggests that embedding the physical law in the loss function provides informative guidance, allowing the PINN to fit the temporal dynamics more intelligently and in a physics-informed manner compared to the FNN_mean_time model. Moreover, although the overall performance of the PINN is only slightly better than that of the FNN model, the PINN is more effective at detecting infected avocadoes by smartly leveraging temporal information in combination with spectral features.

**Figure 6.**
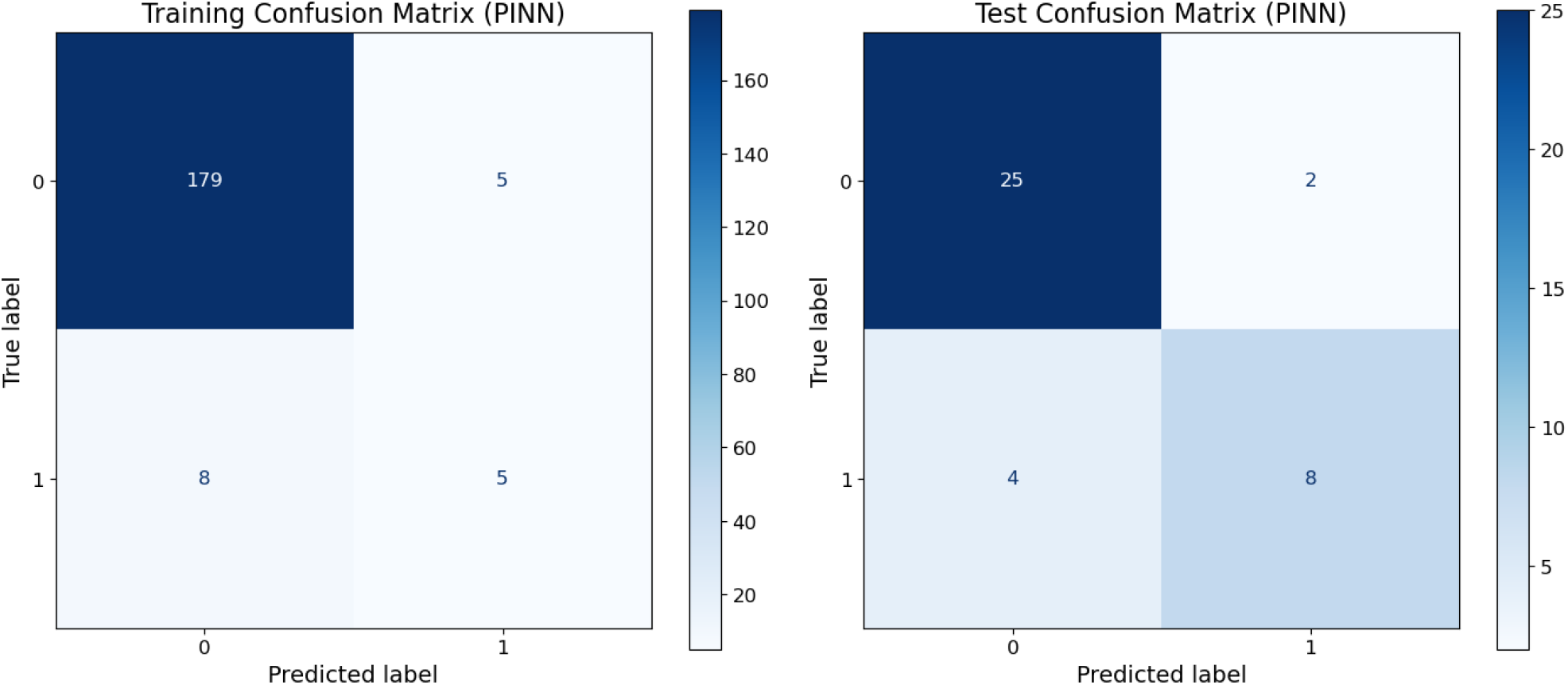
The confusion matrices for the PINN model on the training dataset and the day-12 testing dataset

### 3.2. Results based on random train–test splitting

This section presents the results obtained using a random 80%–20% train–test split. Table 3 reports the overall prediction accuracy of the three models. An interesting observation is that the FNN_mean_only and FNN_mean_time models achieve identical prediction accuracy on both the training and testing datasets. Their confusion matrices are also identical (see Figure 7 and Figure 8), indicating that the inclusion of time as an additional feature does not influence model performance. This can be explained by the nature of the random data split. Since both the training and testing datasets contain samples from all storage durations, their distributions are highly similar. As a result, the testing accuracy is only slightly lower than the training accuracy, reflecting a relatively easy generalization task. In this context, temporal information does not provide additional discriminative power and may even introduce redundancy or noise, limiting its contribution to model performance. This observation is further supported by the performance of the PINN model. The incorporation of the temporal physics-based rule leads to a decrease in performance on both the training and testing datasets, suggesting that the imposed temporal structure may conflict with the spectral information under a random data split. As a result, the interaction between spectral and temporal features becomes less coherent, ultimately reducing the effectiveness of the model.

**Table 3.** Overall model prediction accuracy on the training dataset and the randomly sampled 20% testing dataset.

| Model | Training accuracy | Testing accuracy |
| --- | --- | --- |
| FNN_mean_only | 0.931 | 0.917 |
| FNN_mean_time | 0.931 | 0.917 |
| PINN | 0.920 | 0.896 |

**Figure 7.**
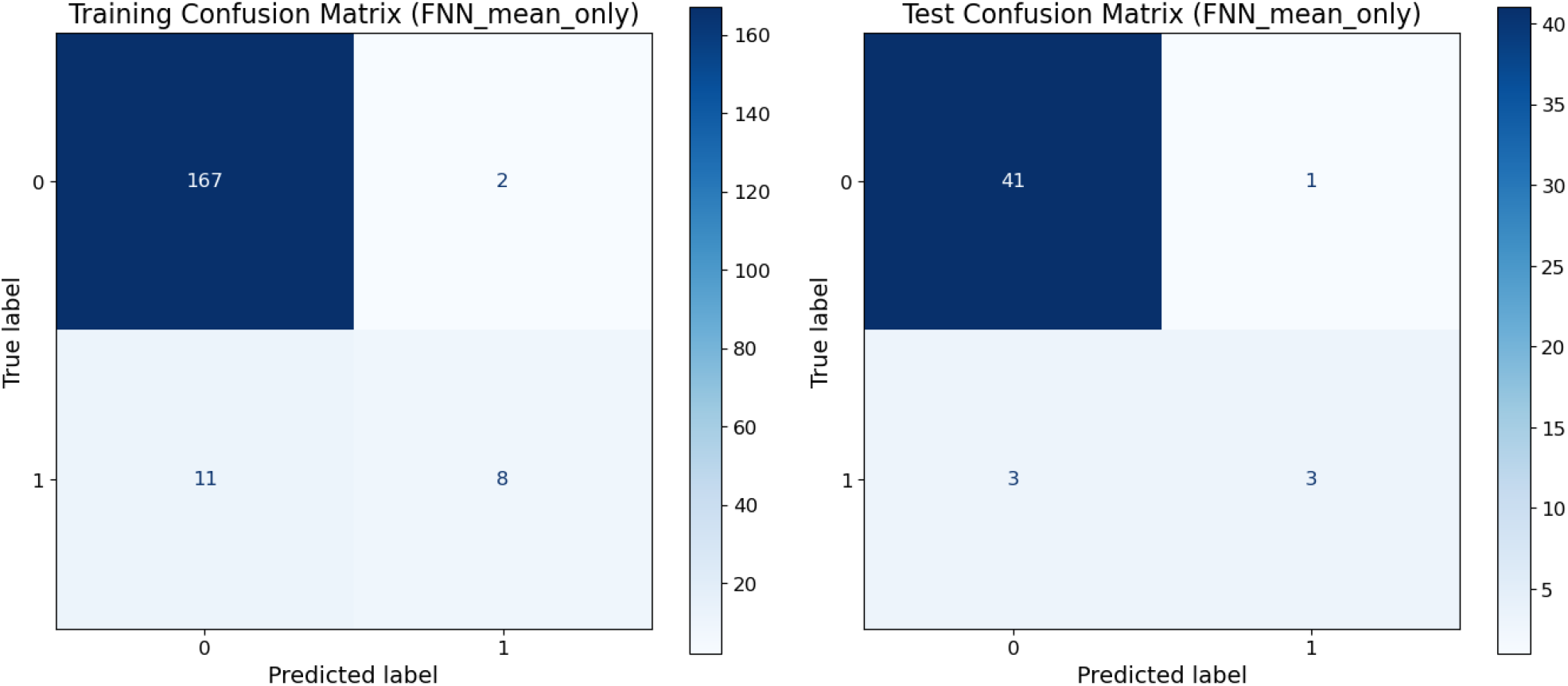
The confusion matrices for the FNN_mean_only model on the training dataset and the randomly sampled 20% testing dataset

**Figure 8.**
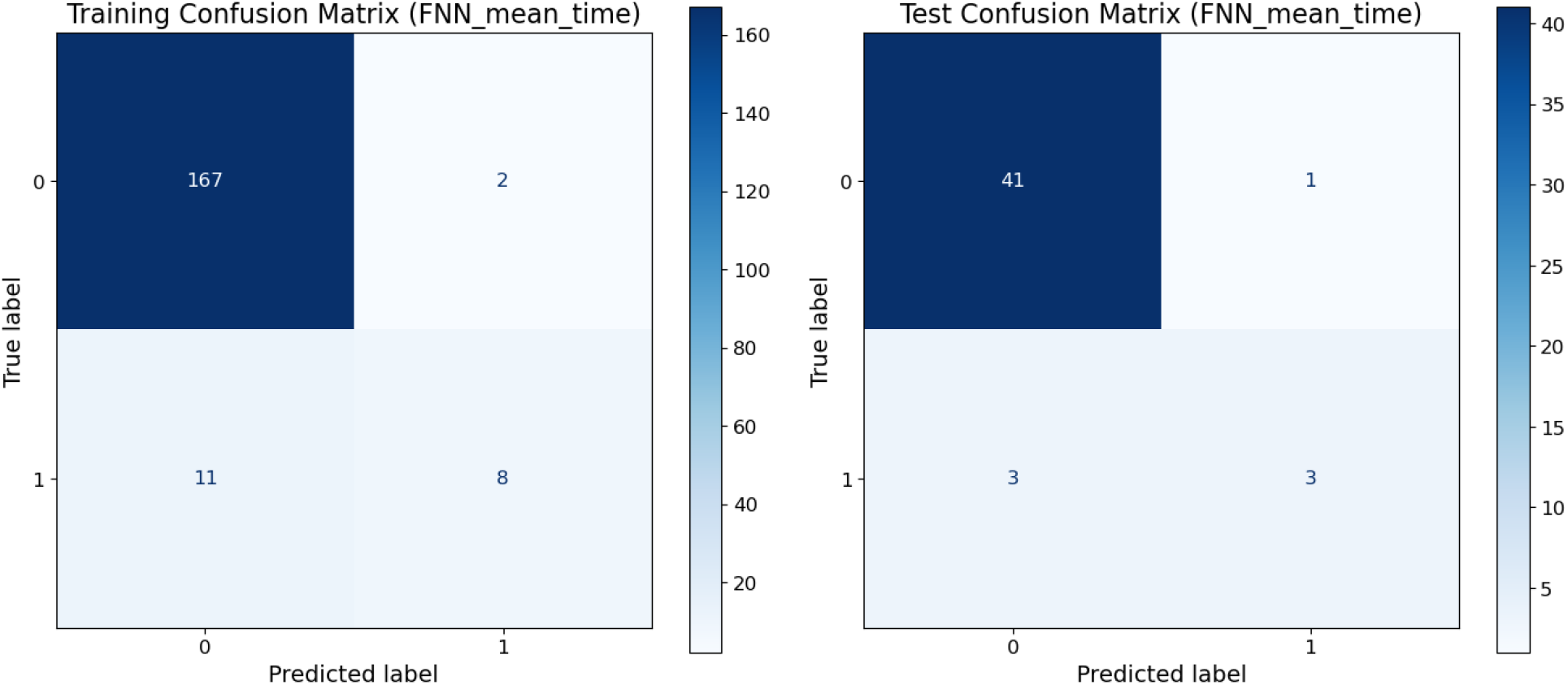
The confusion matrices for the FNN_mean_time model on the training dataset and the randomly sampled 20% testing dataset

**Figure 9.**
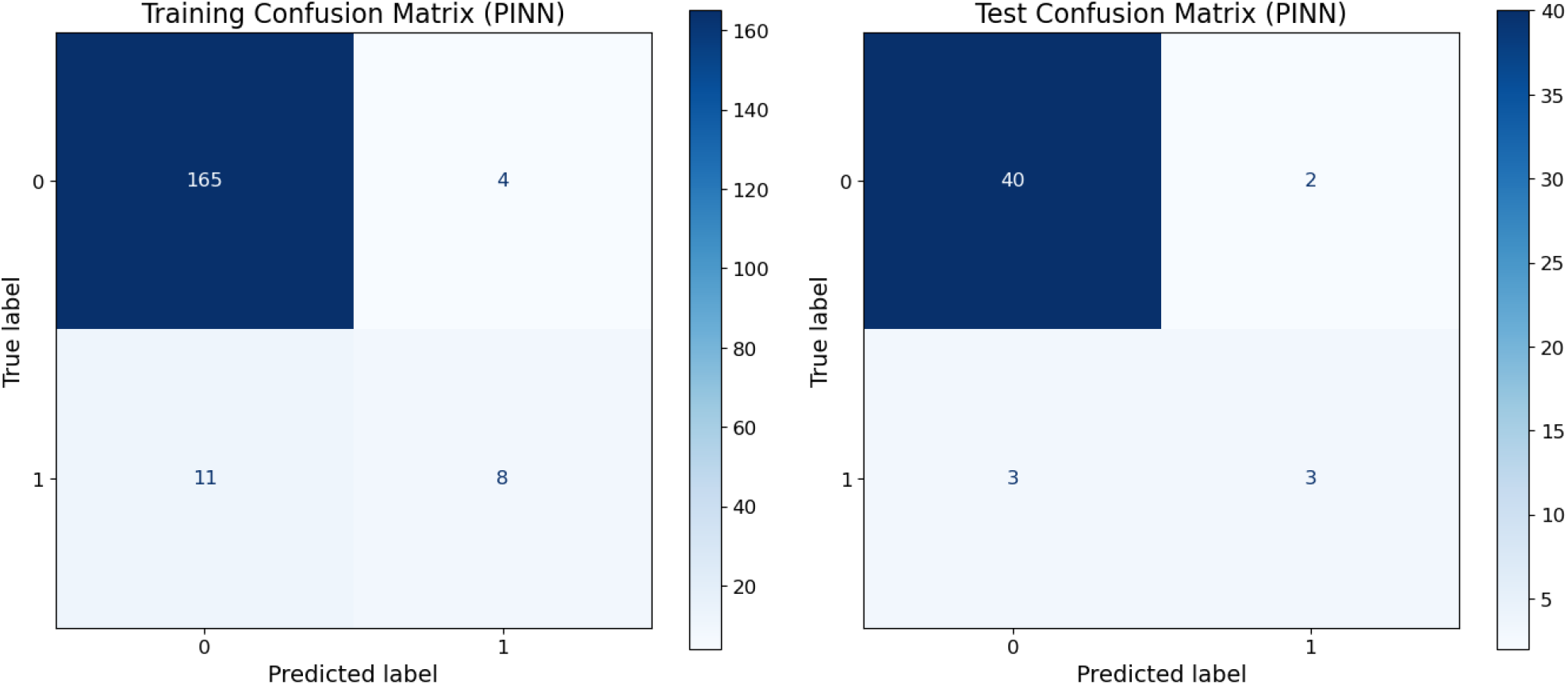
The confusion matrices for the PINN model on the training dataset and the randomly sampled 20% testing dataset

## 4. Conclusion

This study investigated the integration of temporal information into hyperspectral-based machine learning models for predicting avocado stem-end rot during storage. Three modelling approaches were compared: a feedforward neural network using only spectral information (FNN_mean_only), a feedforward neural network incorporating both spectral and temporal information (FNN_mean_time), and a physics-informed neural network (PINN) that embedded a first-order kinetic constraint into the learning process. The results demonstrate that simply adding time as an additional input feature in a purely data-driven manner does not necessarily improve predictive performance and may even increase overfitting to temporal patterns. This was particularly evident from the large number of false positive predictions produced by the FNN_mean_time model when evaluated on the unseen day-12 dataset.

In contrast, the PINN model achieved the highest testing accuracy and substantially reduced false positive classifications while maintaining strong detection capability for infected avocados. These findings suggest that the physics-informed constraint acts as a form of regularization, guiding the neural network to learn temporal dynamics in a biologically meaningful way rather than relying solely on statistical correlations. Importantly, the improvement was most evident under the temporally extrapolated testing scenario, where the models were required to predict infection progression beyond the time range observed during training.

A key contribution of this work is the demonstration that the incorporation of temporal effects through physics-informed learning can help mitigate the sparse-positive problem in food quality datasets. Because infected samples are relatively limited and unevenly distributed across storage durations, purely data-driven models tend to either under-detect infected samples or overcompensate by generating excessive false positives. By embedding mechanistic knowledge about infection growth into the neural network training process, the PINN provides a more balanced utilization of spectral and temporal information, leading to improved generalization and more stable classification behaviour.

Several limitations of the present study should be acknowledged. First, the dataset is relatively small, comprising only 236 avocados with 25 infected samples, which may limit the robustness of the conclusions and increase the uncertainty associated with model estimation. Second, the proposed first-order kinetic formulation represents a simplified description of stem-end rot development. The actual infection process is influenced by multiple biological, physiological, and environmental factors that may not be fully captured by a single kinetic parameter. Third, the dataset was collected under controlled laboratory conditions and from a single experimental setting. Consequently, the developed models may not fully reflect the variability encountered in commercial supply chains, where factors such as temperature fluctuations, handling practices, cultivar differences, and varying infection pressures can affect disease progression. Additional validation under industrial conditions is therefore required before practical deployment.

Despite these limitations, this study highlights the potential value of combining spectral sensing with temporal physics-based modelling in postharvest quality assessment. The findings demonstrate that integrating mechanistic knowledge with machine learning can improve temporal generalization and provide more interpretable predictions than purely data-driven approaches. More broadly, the study draws attention to the importance of incorporating temporal physics-based information into spectral modelling frameworks for food quality prediction. Future research should further validate the proposed approach using larger datasets, multiple storage conditions, different fruit varieties, and more advanced mechanistic formulations, while also exploring its implementation in real industrial monitoring systems.

## Acknowledgement

The authors would like to acknowledge the funding from the Wageningen University & Research “Data Driven & High Tech (DDHT) program (KB-38) with the project number KB-38-001-003” that is sup ported by Dutch Ministry of Agriculture, Fisheries, Food Security and Nature.

## Statement on avocado collection

The avocados were imported from Peru by the project partners (industry partners) under the PPP project Sustainable G&F chains (GreenCHAINge) with TKI ref.: KV1406031.

## Declaration of generative AI and AI-assisted technologies

Generative AI was used to support programming and to improve the clarity, grammar, and readability of the manuscript. All AI-assisted outputs were reviewed and verified by the authors, who take full responsibility for the accuracy, integrity, and final content of the manuscript.

## Note on the preliminary results

The results presented in Section 3.1 are sensitive to the random seeds used. These results should therefore be considered preliminary and will require further investigation.

## Reference

Amoriello, T., Ciorba, R., Ruggiero, G., Masciola, F., Scutaru, D., & Ciccoritti, R. (2025). Vis/NIR spectroscopy and Vis/NIR hyperspectral imaging for non-destructive monitoring of apricot fruit internal quality with machine learning. Foods, 14(2), 196.

Boyce, W. E., DiPrima, R. C., & Brannan, J. R. (2009). Elementary differential equations (Vol. 9). Wiley New York.

Chauhan, A., de Villiers, H., Meesters, L., Paillart, M., Grbović, Ž., Panić, M., & Brdar, S. (2024). Avocado stem-end rot detection using hyperspectral imaging. VII International Conference Postharvest Unlimited,

Chauhan, A., Hemming, S., & Nicolaï, B. M. (2023). AI in supply chain management of horticultural products. In Artificial intelligence in the agri-food sector: Applications, risks and impacts (pp. 45–48).

Elsayed, S., Gala, H., Abd El-baki, M. S., Maher, M., Elbeltagi, A., Salem, A., Elwakeel, A. E., Elsherbiny, O., & Abd El-Fattah, N. G. (2025). Hyperspectral technology and machine learning models to estimate the fruit quality parameters of mango and strawberry crops. Plos one, 20(2), e0313397.

Guo, X., Harbers, C. J., Heuer, H. E., Min, Q., Liu, Y., Woltering, E. J., Marcelis, L. F., & van der Sman, R. G. (2026). A generic scientific machine learning framework for fruit and vegetable quality prediction. Postharvest Biology and Technology, 234, 114103.

Lammertyn, J., Aerts, M., Verlinden, B. E., Schotsmans, W., & Nicolaı, B. M. (2000). Logistic regression analysis of factors influencing core breakdown in ‘Conference’pears. Postharvest Biology and Technology, 20(1), 25–37.

Lamptey, F. P., Teye, E., Abano, E. E., & Amuah, C. L. (2023). Application of handheld NIR spectrometer for simultaneous identification and quantification of quality parameters in intact mango fruits. Smart Agricultural Technology, 6, 100357.

Mancini, M., Mazzoni, L., Gagliardi, F., Balducci, F., Duca, D., Toscano, G., Mezzetti, B., & Capocasa, F. (2020). Application of the non-destructive NIR technique for the evaluation of strawberry fruits quality parameters. Foods, 9(4), 441.

Singh, R., Ruhil, A., Jain, D., Patel, A., & Patil, G. (2009). Prediction of sensory quality of UHT milk–a comparison of kinetic and neural network approaches. Journal of food engineering, 92(2), 146–151.

Téllez-García, J., Sánchez, M.-T., González-Fernández, J.-J., & Pérez-Marín, D. (2025). Evaluation of NIRS portable sensors for the determination of the physicochemical quality and maturity of intact mangoes. Scientia Horticulturae, 354, 114514.

Tijskens, L., & Polderdijk, J. J. (1996). A generic model for keeping quality of vegetable produce during storage and distribution. Agricultural Systems, 51(4), 431–452.

Tromp, S.-O., Haijema, R., Rijgersberg, H., & van der Vorst, J. G. (2016). A systematic approach to preventing chilled-food waste at the retail outlet. International Journal of Production Economics, 182, 508–518.

Tromp, S.-O., Harkema, H., Hogeveen, E., Rijgersberg, H., & Woltering, E. J. (2017). On the validation of improved quality-decay models of potted plants. Postharvest Biology and Technology, 123, 119–127.

Tromp, S.-O., Rijgersberg, H., da Silva, F. P., & Bartels, P. (2012). Retail benefits of dynamic expiry dates—Simulating opportunity losses due to product loss, discount policy and out of stock. International Journal of Production Economics, 139(1), 14–21.

van der Sman, R., Guo, X., & Pedreschi, R. (2026). Physics-Informed Neural Network for prediction of post-harvest firmness of avocados. Postharvest Biology and Technology, 231, 113898.

Wang HaiLong, W. H., Peng JiYu, P. J., Xie ChuanQi, X. C., Bao YiDan, B. Y., & He Yong, H. Y. (2015). Fruit quality evaluation using spectroscopy technology: a review.

Xiao, X., & Li, M. (2022). Fusion of data-driven model and mechanistic model for kiwifruit flesh firmness prediction. Computers and electronics in agriculture, 193, 106651.

Zhu, H., Chu, B., Fan, Y., Tao, X., Yin, W., & He, Y. (2017). Hyperspectral imaging for predicting the internal quality of kiwifruits based on variable selection algorithms and chemometric models. Scientific Reports, 7(1), 7845.

